## Supplementary material for "A shared neural code for the physics of actions and object events": Supplemantary Text

**Supplementary Information**

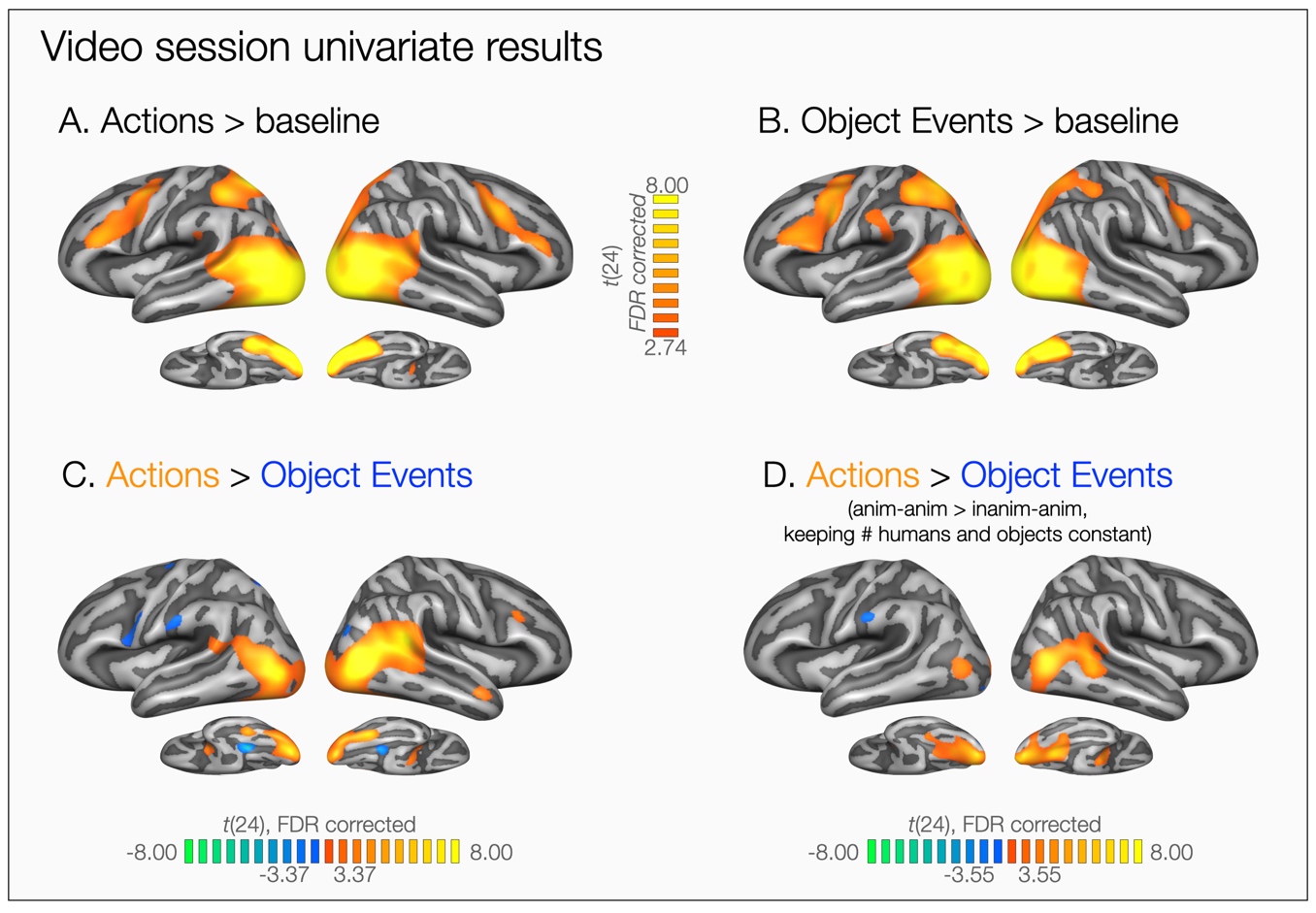

**Supplementary Figure 1 – Video session univariate results.** Whole-brain maps showing univariate changes in neural responses to actions and object events. All maps are FDR-corrected. (**A-B)** Compared against the baseline, both actions and object events led to increased signal in bilateral posterior occipitotemporal regions, parietal lobes, and premotor cortex. (**C)** Whole-brain contrast of all actions versus all object events. Stronger activity is observed in posterior superior temporal sulcus and lateral occipitotemporal cortex for actions compared to object events, particularly in the right hemisphere. In ventral occipitotemporal cortex, stronger responses to actions are observed more laterally, while stronger responses to object events are observed more medially. This gradient is consistent with the known organization of ventral occipitotemporal cortex for animate versus inanimate object categories^1^. Note that some of the differences in this contrast can be attributed to the number of bodies and objects in the scene. (**D)** Whole-brain contrast of animate-inanimate (e.g., the boy jumps over the box) > inanimate-animate (e.g., the ball bounces over the girl) events. This contrast balances for the number of bodies and objects in the scene. Keeping the number of humans and objects in the scene constant, stronger activity is observed for actions compared to object events in bilateral ventral occipitotemporal cortices, and lateral occipitotemporal cortex, particularly in the right hemisphere.

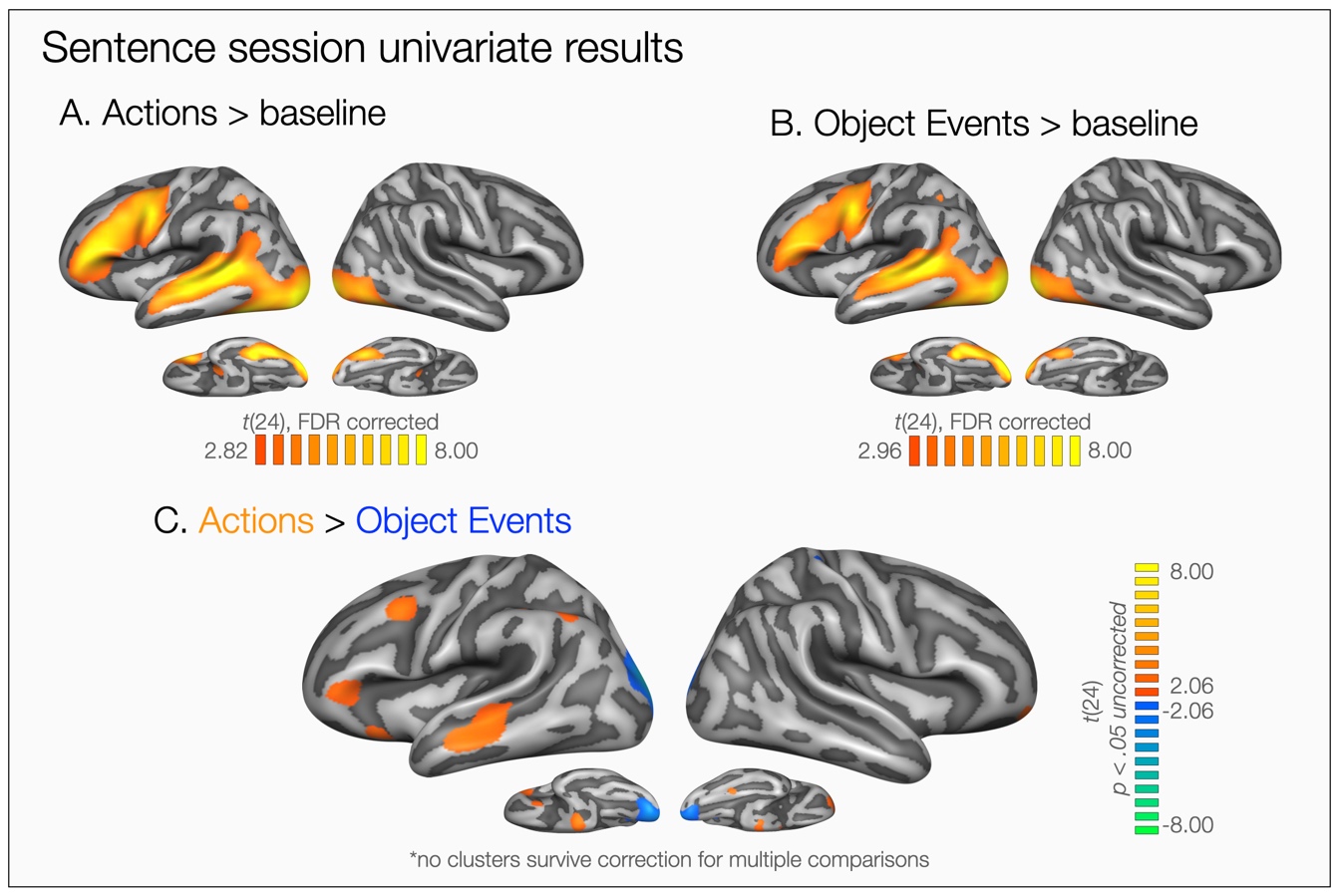

**Supplementary Figure 2 – Sentence session univariate results. (A-B)** Whole-brain maps showing univariate changes in neural responses to sentences describing actions and object events. Both maps are FDR-corrected. Overall, compared against the baseline, both actions and object event sentences led to increased signals in the left hemisphere particularly in temporal gyrus and sulcus, inferior parietal lobe, and premotor cortices. **(C)** Uncorrected whole-brain contrast of actions and object events in the sentence session. The map is thresholded at *p* < .05 to demonstrate differences that do not survive FDR-correction. No clusters survived correction for multiple comparisons.

**Supplementary Note 1 - Decoding of actions and object events separated by animate and inanimate patients.**

Previous work revealed different neural responses to human actions with or without human recipients in several brain regions, especially in the right superior temporal sulcus^2^. Our design allowed us to address whether these effects reflect ‘sociality’ of the action or the animacy of the affected entity. To address this question, we investigated decoding accuracies for actions and object events separated by patient type: animate-animate & animate-inanimate, inanimate-animate & inanimate-inanimate events (Supplementary Figures 3a-b). Overall, despite differences in decoding strength, we observed above chance decoding of the three events in frontoparietal and posterior temporal cortices linked to action recognition. Since previous work suggested right superior temporal sulcus as a key region for encoding social interactions, we also compared decoding strengths for animate (in orange) and inanimate recipients (in green) for actions and object events in right and left STS/pMTG (Supplementary Figure 3c).

A linear mixed effects model revealed an event type by target interaction in both left (χ2[1] = 9.54, *p* = .002, ΔAIC = 7.54) and right hemispheres (χ2[1] = 4.46, *p* = .035, ΔAIC = 2.46). In both left and right pSTS, we observed better decoding for actions with animate versus inanimate patients (‘social’ versus ‘non-social’) (Left - *b* = 7.61, *t*(72) = 3.08, *p* = .003, *d* = .87, 95% CI [.29 1.45]; Right - *b* = 5.14, *t*(72) = 2.17, *p* = .033, *d* = .62 , 95% CI [.04 1.19]), but not for object events (Left - *b* = -.47, *t*(72) = -.19, *p* = .851, *d* = -.05, 95% CI [-.62 .51]; Right - *b* = 3.61, *t*(72) = 1.53, *p* = .131, *d* = .43 , 95% CI [-.14 1.00]). This pattern fits nicely with the previous literature and shows that in STS/pMTG, the animacy of the patient is relevant only for animate subjects. We would like to note that we found this effect in both right and left pSTS. Future work should address what aspects of 'social’ versus ‘non-social’ actions are represented in the left and right posterior temporal cortices.

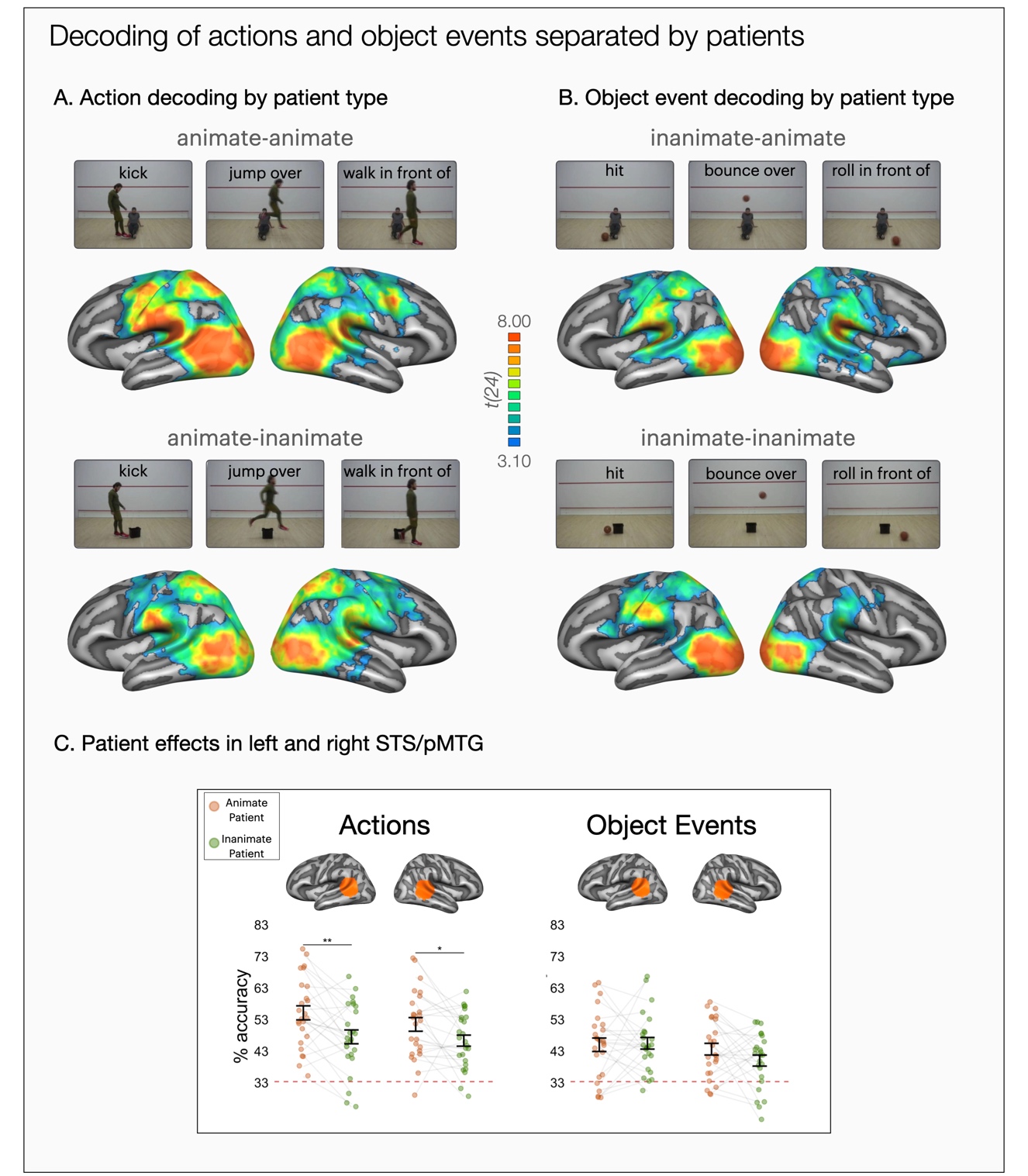

**Supplementary Figure 3** *-* **Decoding of actions and object events separated by animate and inanimate patients.** Decoding accuracies for actions and object events separated by patient type: **(A)** animate-animate & animate-inanimate, **(B)** inanimate-animate & inanimate-inanimate events. The maps were thresholded by areas corrected for multiple comparisons using Monte Carlo Cluster based correction (*p*_initial_ = .001). **(C)** FDR-corrected pairwise two-tailed tests of estimated marginal means for actions and objects with animate (in orange) or inanimate recipients (in green) in right and left STS/pMTG (* *p* < .05, ***p* < 0.01, ****p* < 0.001). Error bars indicate standard error of the mean (SEM, n=25).

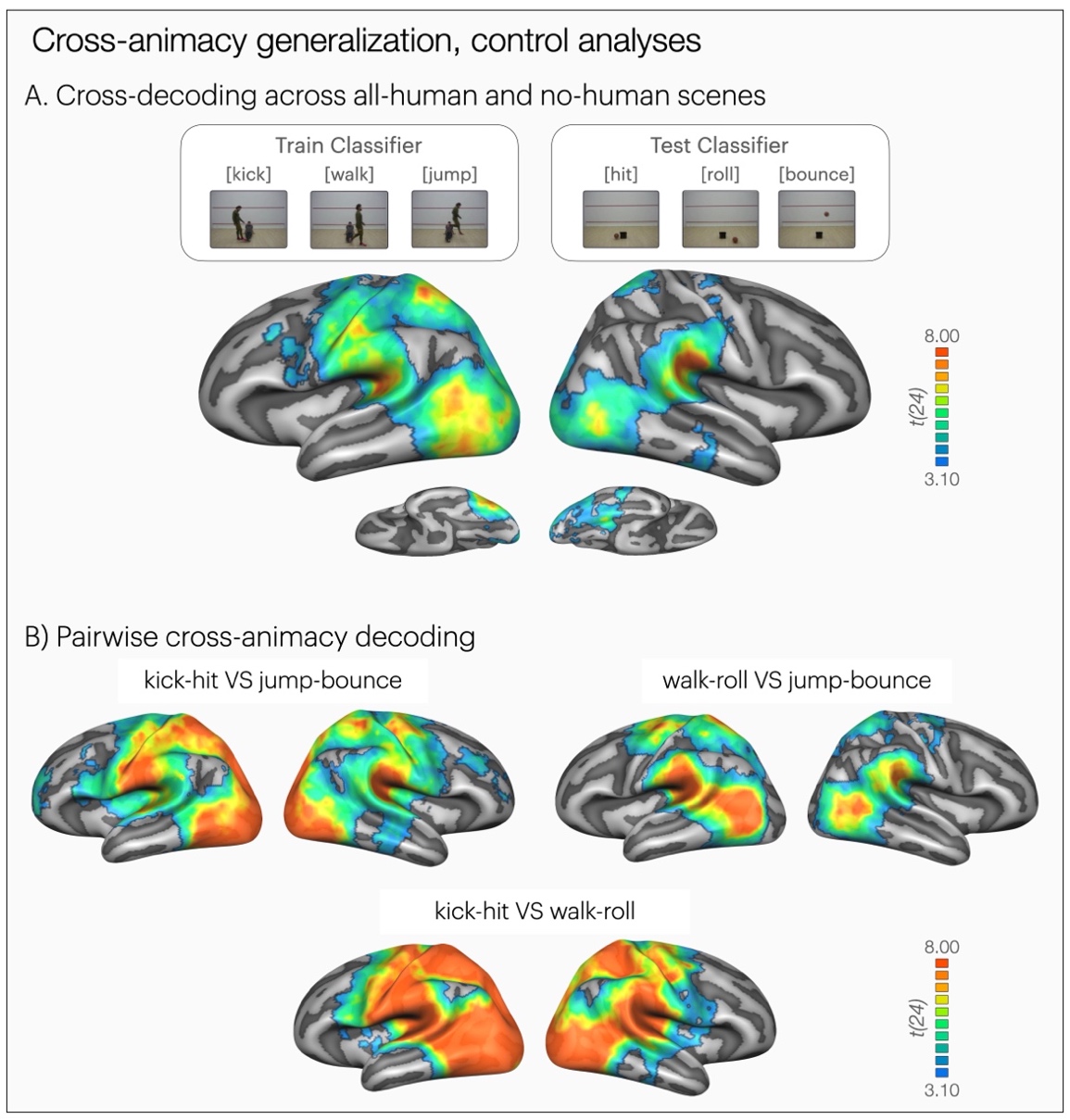

**Supplementary Figure 4** *-* **Control analyses for cross-animacy generalization of observed events.** Results of whole-brain three-way (**A)** cross-decoding of actions and object events across all-human and all-object scenes (e.g., train on ‘*the boy jumps over the man’* test on ‘*the ball bounces over the box’*). One-tailed t-tests against chance level 33.33%. The maps are thresholded by areas corrected for multiple comparisons using Monte Carlo Cluster based correction (*p*_initial_ = .001). Even though the effects are slightly weaker compared to the cross-decoding where all conditions were included in the analysis, we still observe above chance decoding in premotor cortex, inferior parietal lobe, and posterior temporal lobes. This suggests that the cross-decoding results cannot be explained away by the relevance of some object events for animate entities in the scene since cross-decoding persists even when the object events do not pertain to an animate being. Note that for this analysis, the data used for decoding decreases in half, which could partly explain the overall decrease in decoding strength. **(B)** Results of whole-brain three-way decoding searchlight for cross-decoding of actions and object events to test if successful cross-decoding relies on a peculiar event being highly distinct from the others. One-tailed t-tests against chance level 50%. The maps are thresholded by areas corrected for multiple comparisons using Monte Carlo Cluster based correction (*p*_initial_ = .001). Based on these maps, we can suggest with confidence that cross-decoding in frontoparietal and posterior temporal cortices does not rely on one peculiar event since cross-animacy decoding for pairs of motion events revealed qualitatively similar patterns of decoding across the core regions of the action observation network.

**Supplementary Note 2** *-* **Whole-brain GLM representational similarity analyses testing the representation of inter-object relations and motion path***.*

What shared aspects of actions and object events are captured by cross-animacy decoding? As a first step at characterizing the representational organization of brain regions with respect to events, we built two models of representational similarity specifying different event components: inter-object relations and motion trajectory. For inter-object relations, we built a binary model specifying presence/absence of contact in the stimulus set (i.e., hit-kick events involved contact between two objects, while jump-bounce and walk-roll events did not). For motion path, we built a binary model specifying horizontal versus vertical movement in the depicted scenes (i.e., hit-kick and roll-walk events included horizontal motion only, while jump-bounce events also included vertical motion). We conducted two whole-brain searchlight-based representational similarity analyses using representational dissimilarity matrices obtained from the contact and motion trajectory models.

The RSA for making contact (see Supplementary Figure 5a) revealed significant effects across frontoparietal and posterior temporal cortices that have also shown cross-animacy decoding of observed events (compare to Figure 3). Among the regions of the action observation network, the left anterior IPL showed the strongest effect. Yet, since the presence/absence of contact correlated with other visual characteristics in the scenes, this evidence alone does not guarantee that these shared event representations are not tied to visual cues. The RSA for motion path (i.e., horizontal vs. vertical motion), on the other hand, revealed significant effects only in bilateral occipital poles and bilateral medial premotor cortices, but not in regions of the action observation network (see Supplementary Figure 5b). Overall, these initial findings suggest that frontoparietal and posterior temporal regions linked to action recognition may encode events at a level specifying inter-object relations.

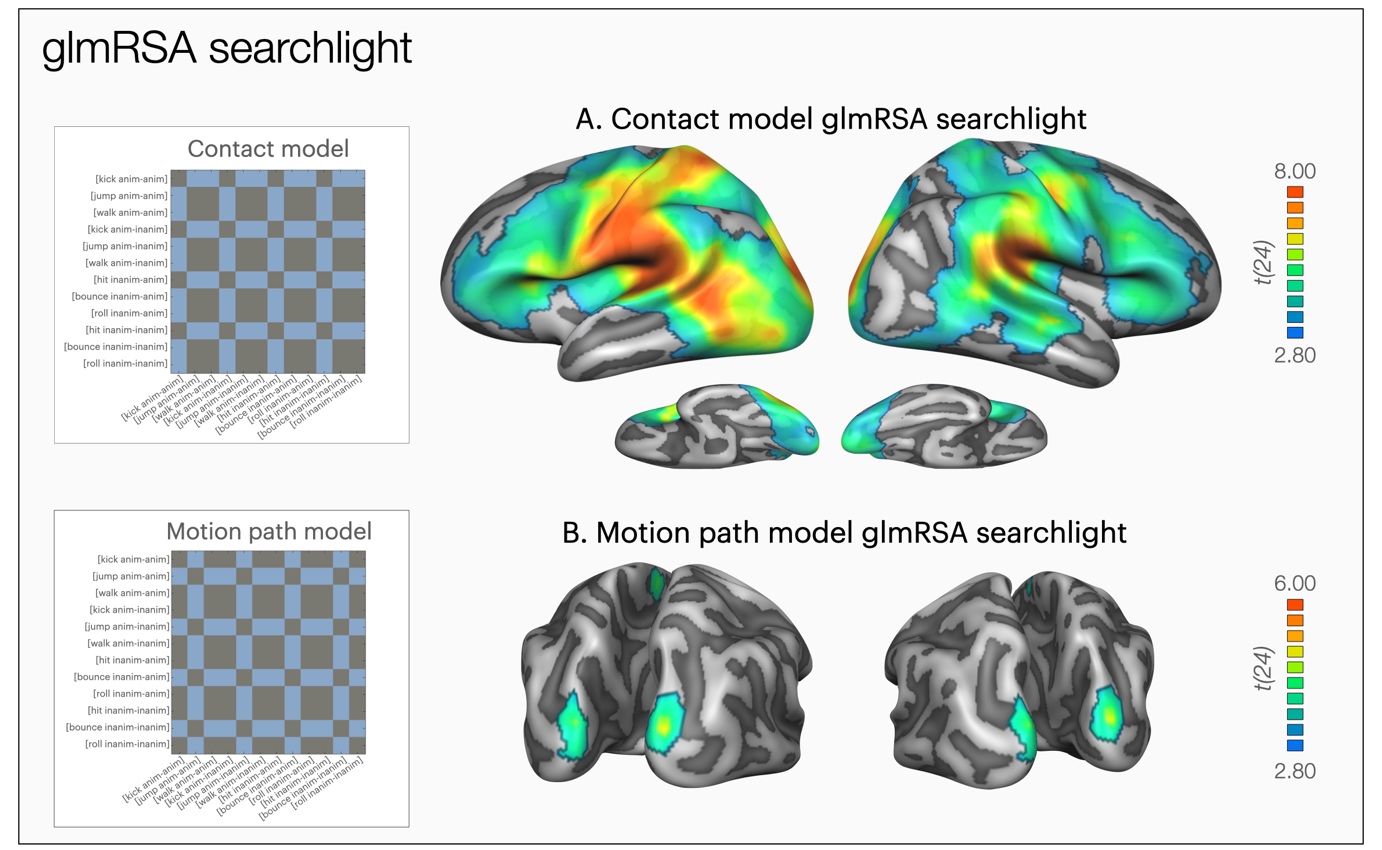

**Supplementary Figure 5 - Whole-brain GLM representational similarity analyses testing the representation of inter-object relations and motion path***.* Whole-brain GLM RSA searchlight for models of **(A)**  making contact (i.e., hit vs. no hit) and **(B)** motion path (i.e., horizontal vs. vertical motion). The maps are thresholded by areas corrected for multiple comparisons using Monte Carlo Cluster based correction (*p*_initial_ = .001).

**Supplementary Note 3 - Within-sentence and cross-modal decoding of events**

In the sentence session, action and object events were decoded above chance in all left hemisphere ROIs (see Supplementary Figure 6e). Cross-modal decoding of actions was successful in LOTC bilaterally, as well as in pSTS and angular gyrus (see Supplementary Figure 6c-f). Cross-modal decoding of object events did not yield very robust and reliable results in the whole-brain or ROI analyses (see Supplementary Figure 6d-f). Cross-modal decoding of object events was also weaker compared to cross-animacy + modality generalization (compare Supplemental Figure 6d to Figure 4b). This could be surprising given the fact that cross-animacy + modality decoding requires two steps of generalization while cross-modal decoding of object events requires only one step of generalization. However, comparing the decoding strength across these decoding schemes is not straightforward since cross-modality decoding of object events relies on video and sentence data of object events only (50% of the data, as opposed to 100% of the data for cross-animacy + modality decoding). Furthermore, the decoding of object event sentences and cross-modal decoding of object events were generally weaker compared to that of actions. However, cross-animacy + modality decoding used the entire dataset that also benefits from stronger results for human action sentences.

There could be various reasons for the difference between decoding strength for action and object events, both in sentences and across modality. First, the verbs or subjects we used for object event sentences might have resulted in more variable neural activity patterns compared to verbs that were used for action sentences. This might have contributed to weaker cross-modal decoding for object events as well. Furthermore, the weak cross-modal decoding for object events compared to actions might imply that the visual-to-verbal reference is clearer and more specific in the context of actions, and more variable in the context of object events. Future studies with sentence stimuli that are more controlled in terms of their formal and visual properties could address these issues.

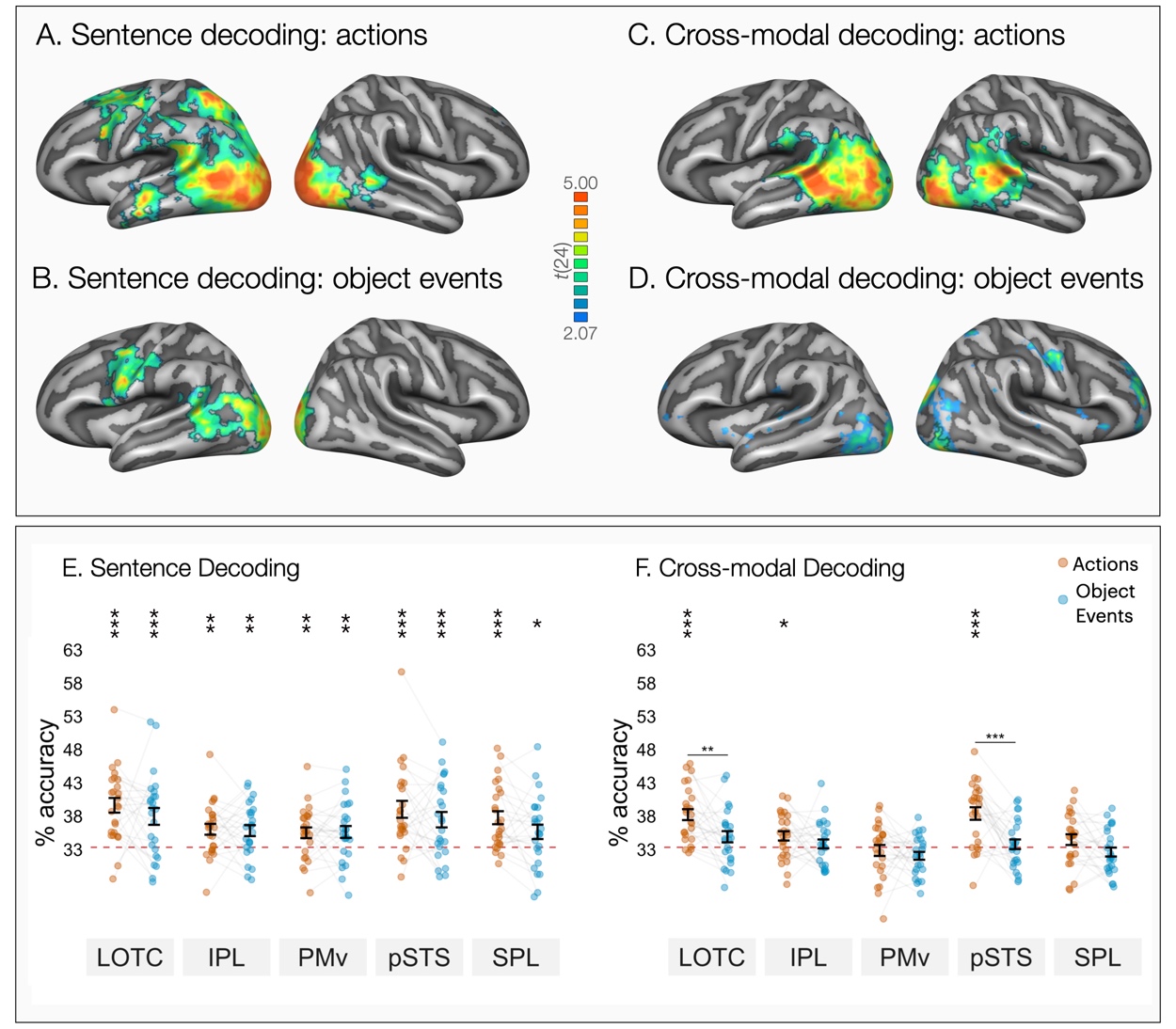

**Supplementary Figure 6 - Within-sentence and cross-modal decoding of events.** Whole-brain three-way decoding of **(A)** action sentences, **(B)** object event sentences, **(C)** actions across modality, and **(D)** object events across modality, (one-tailed t-tests against chance level of 33.33%). Maps in A-C are thresholded for areas that survive correction for multiple comparisons using Monte Carlo Cluster based correction (*p*_initial_ = .005). The map in D (cross-modal decoding of object events) is thresholded at *p* < .05 to demonstrate trends that do not survive correction. For action and object event decoding across modality (C-D), a classifier was trained on videos of one event type (e.g., action videos) and tested on sentences of the same event type (e.g., action sentences). **(E)** Left hemisphere ROI decoding accuracies for action and object event sentences. **(F)** Left hemisphere ROI decoding accuracies for actions and object events across modality. Error bars in (E-F) indicate standard error of the mean (SEM, n=25) and asterisks indicate FDR-corrected effects of one-tailed t-tests for comparisons against chance-level (33.33%, * *p* < .05, ***p* < 0.01, ****p* < 0.001). FDR-corrected two-tailed estimated marginal means analyses showed better cross-modal decoding for actions than object events in left LOTC (*b* = 3.32, *t*(216) = 3.27, *p* = .001, *d* = .93 , 95% CI [.36 1.49]) and pSTS (*b* = 4.61, *t*(216) = 4.54, *p* < .001, *d* = 1.29 , 95% CI [.72 1.85]).

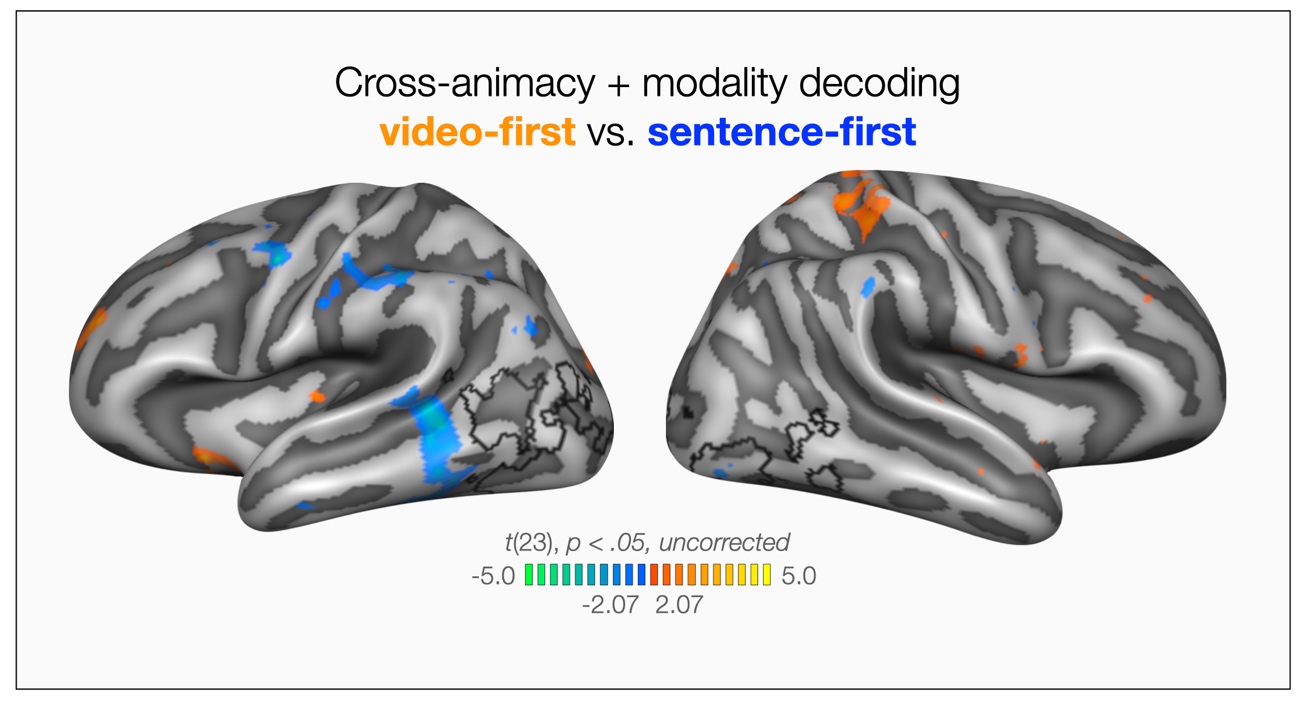

**Supplementary Figure 7** *-* **Modulation of cross-animacy + modality decoding accuracies by session order.** It is possible that cross-animacy + modality generalization in left LOTC was due to verbalization (in the video session) or visual imagery (in the sentence session). If across modality generalization is due to imagery, we would expect stronger imagery for people who received the videos first and sentences second, and stronger verbalization for people who received sentences first and videos second. The figure displays the output of a two-tailed independent samples t-test between decoding accuracies of the participant group who completed the sentences first (in blue) and the participant group who completed the videos first (in orange). To demonstrate any trends of session order effects, maps were thresholded at *p* < .05. The black outline marks the areas that showed above-chance cross-animacy + modality decoding as reference (compare to Figure 4b). Contrasting the decoding accuracies of the two groups revealed no significant differences indicative of an effect of imagery or verbalization. Thus, we found no support for the hypothesis that visual imagery and verbalization can fully account for the observed generalization across animacy and modality.
